# TRICLOSAN IS A KCNQ3 POTASSIUM CHANNEL ACTIVATOR

**DOI:** 10.1101/2021.09.30.462664

**Authors:** Victor De la Rosa, Maria Luisa Guzmán-Hernández, Elisa Carrillo

## Abstract

KCNQ channels participate in the physiology of several cell types. In neurons of the central nervous system, the primary subunits are KCNQ2, 3 and 5. Activation of these channels silence the neurons, limiting action potential duration and preventing high-frequency action potential burst. Mutation of the KCNQ genes are associated with a wide spectrum of phenotypes characterized by hyperexcitability. Activation of KCNQ channels is an attractive strategy to treat epilepsy and other hyperexcitability conditions as are the evolution of stroke and traumatic brain injury. In this work we show that triclosan, a bactericide widely used in personal care products, activates the KCNQ3 channels but not the KCNQ2. Triclosan induces a voltage shift in the activation, increases the conductance and slows the closing of the channel. The effect is independent of PIP2. The putative binding site is located in the pore region but is distinct from the binding site for retigabine. Our results indicate that triclosan is a new activator for KCNQ channels.

## Introduction

KCNQ (Kv7) potassium channels play an important role in the control of membrane potential and neuronal excitability [1]. Except for KCNQ1 which is found predominantly in the heart, KCNQ2-5 channels are mainly neuronal. These channels generate a muscarinic inhibited potassium current that controls excitability, referred to as the M-current, which is controlled by membrane voltage and PIP2. Activation of neuronal KCNQ channels by small molecules has emerged as a therapeutic strategy for treatment of hyperexcitability related disorders [2, 3]. For example, Retigabine (RTG) is a widely used antiepileptic drug that shifts to negative potentials the voltage dependence of activation of KCNQ2/3 channels (heteromeric channels with KCNQ2 and KCNQ3 subunits) [4]. This shift allows the channels to be open at potentials similar to basal membrane values, this facilitation shuts down cell hyperexcitability observed in anxiety, mania, tinnitus, epilepsy, stroke and traumatic brain injury [3]. Although significant advances have been made in recent years, the search for KCNQ channel openers is still under intensive investigation since it can help design new selective modulators to target disorders more precisely. The molecular determinants of action of different drugs have been a subject of recent studies. For example, ICA-27243, a KCNQ opener, retained activity on channels lacking the conserved tryptophan in the pore region of the channel that is essential for RTG activity. Chimeric channels revealed that ICA-27243 binding site is located on the VSD [5], indicating that different drugs have distinct binding sites and that the mechanism of action may be different. This heterogeneity might be the key to find drugs with different KCNQ channels subtype specificity.

Originally developed as a pesticide, triclosan was used primarily as a bactericide found in many personal care products [6] such as toothpaste and soaps. In 2016, the United States Food and Drug Administration banned triclosan in antiseptic wash products. However, triclosan persist in the environment, in fact the disposal of these products led to detectable levels of triclosan in treated wastewater, ocean water and fresh water [7–9]. Triclosan is hydrophobic and can accumulate in wild animals, detectable levels are also found in human samples such as urine and breast milk [10–12].

Exposure to environmental levels of triclosan may affect human health via different mechanisms. Triclosan interacts with the ryanodine receptors RyR1 and RyR2, thus disturbing the excitation-contraction coupling and Ca^2+^ dynamics in striated muscle and L-type channels Ca^2+^ entry in cardiac muscle [13, 14]. In mast cells, triclosan caused mitochondrial fission by reactive oxygen species (ROS) [15]. There is evidence that triclosan at induces hyperpolarization of the membrane potential of lymphocytes, presumably due to activation of Ca^2+^ dependent K^+^ channels [16]. Triclosan has also been shown to produce toxic effects in neural stem cells via ROS [17], and to decrease the expression of N-methyl-D-aspartate subunits, leading to evidence that triclosan acts as a neurotoxic agent. Moreover, it has been shown that triclosan perturbs the hippocampal synaptic plasticity of rats [18].

To our knowledge, no studies have been undertaken to investigate the effects of triclosan directly on potassium channels. In this study, we show that triclosan selectively activates the KCNQ3 channel. Mutagenesis analyses suggest that the binding site of triclosan is different from that of retigabine and other voltage sensing domain binding drugs. Triclosan can be used as a tool for investigation of new synthesized compounds that bind selectively to KCNQ channels members.

## Experimental procedures

### Cell culture and transfection

HEK-293 cells were grown on tissue culture dishes in Dulbecco’s modified Eagle’s medium (Thermo Fisher Scientific) with 10% heat-inactivated fetal bovine serum (Gibco) plus 1% penicillin/streptomycin in a humidified incubator at 37 °C and 5% CO_2_ and passaged every 3-4 days using 0.05% of Trypsin-EDTA (Thermo Fisher Scientific) to lift the cells from the culture dish. For transfection, cells were plated into 24 well plastic culture dishes and transfected 24 hours later with Lipofectamine 2000 (Thermo Fisher Scientific) according to manufacturer’s instructions. The next day cells were plated onto coverslips and experiments were performed over the following 1 – 2 days. Cells were cotransfected with the indicated cDNA, together with EGFP cDNA (pEGFP) as a transfection reporter.

### Molecular biology

The expression plasmid of the KCNQ channels genes is pcDNA3.1. Site directed mutagenesis was performed using the Phusion High-Fidelity PCR master Mix (New England Biolabs). All mutations were confirmed by DNA sequencing.

### Electrophysiology

Current recordings were performed in whole – cell configuration. Pipettes were pulled from borosilicate glass capillaries (BF150-86-10HP; Sutter Instruments) by using a Flaming/brown micropipette puller P-97 (Sutter Instruments), tips were fire polished with the use of a microforge (Narishige). Pipettes had resistances of 2-3 MΩ when filled with internal solution and measured in standard bath solution. Membrane current was measured with an Axopatch 200A patch clamp amplifier and digitized for storage and analysis using a Digidata 1322A interface controlled by pClamp9.2. Series resistance was routinely compensated 60%, liquid junction potential corrections were not applied.

The external Ringer’s solution contained (in mM): NaCl 145, KCl 5, CaCl_2_ 2, MgCl_2_ 1, HEPES 10; pH 7.4. The pipette solution contained (in mM): KCl 140, MgCl_2_ 2, EGTA 10, HEPES 10, GTP 2, ATP-Na_2_ 0.3, phosphocreatine 10; pH 7.3. Triclosan (Sigma) was diluted in ethanol and stocked at 100 mM concentration. Further dilutions were made on Ringer’s solution to desired concentrations. Solutions were delivered by gravity from several reservoirs selectable by activation of solenoid pinch valves (Cole – Parmer). Bath solution exchange was essentially complete by ~30 s. Experiments were performed at room temperature.

To estimate voltage dependence, tail current amplitudes (I_TAIL_) were measured at −20 mV or otherwise stated after a 1000 ms prepulse to depolarizing voltages, normalized, and plotted as a function of test potential. I_TAIL_-V relations were fitted to a Boltzmann function of the form:

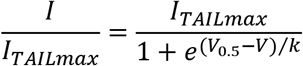

Where I_TAILmax_ is the maximum tail current, V_0.5_ is the voltage that produces half-maximal activation of the conductance and *k* is the slope factor. V_0.5_ values were plotted as a function of Triclosan concentration and fit to a Hill function of the form:

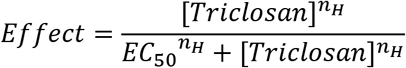

Where EC_50_ is the concentration at which 50% effect is reached and nH is the Hill coefficient. Data shown represents mean ± standard deviation for *n* determinations. Clampfit 10.6 and OriginPro 8.6 software were used for data analysis.

### Homology modelling and docking

The human KCNQ3 channel homology model was created using the iTasser server [19–21] using the KCNQ2 cryo-EM structure as a template (PDBID: 7CR0) [22]. The model is limited to the transmembrane domains. The A315T pore mutation was omitted from the model. The resulting homology model was manually inspected and refined with ModRefiner server [23]. Further structural models were generated by rearrangement of four subunit models as a tetramer. The possible binding sites of the ligand was determined with the AutodockTools 1.5.6 software [24] and PatchDock [25]. One simulated triclosan molecule was simulated docked per subunit. A r.m.s tolerance of 2 Å was used for clustering in AutoDock, and 1.5 Å for PatchDock using a protein-small ligand docking algorithm. The lowest energy docked conformation is shown. Figures were prepared using Pymol 2.3.3 [26–28]

## Results

### Activation of KCNQ2/3 channels by triclosan

The bactericide triclosan was evaluated on the KCNQ2/3 channel. Whole-cell potassium currents from transfected KEH-293 cells are shown in figure 1A. Cells were held at −80 mV and tail currents were measured at −20 mV after consecutive voltage steps ranging from −120 mV to +10 mV. Triclosan shifts the open probability of the channel to negative voltages in a dose – dependent manner (Figure 1B). The maximum voltage shift was ~20 mV. The data was fit to a Hill equation (Figure 1C), giving an EC_50_ = 32 μM and n_H_ = 2.02. This contrast with the voltage shift of the antiepileptic drug Retigabine which is ~50 mV at saturating concentrations (Figure 1D-F). It is worth noting that the nH value is almost double the nH for Retigabine, which might indicate a different mechanism of action. The tail current in the retigabine recordings was measured at −120 mV. Both triclosan and retigabine at saturating concentrations slow the tail current kinetics by a factor of 2.5 and 3.4 respectively (measured at −100 and −120 mV correspondingly).

**Figure 1.**
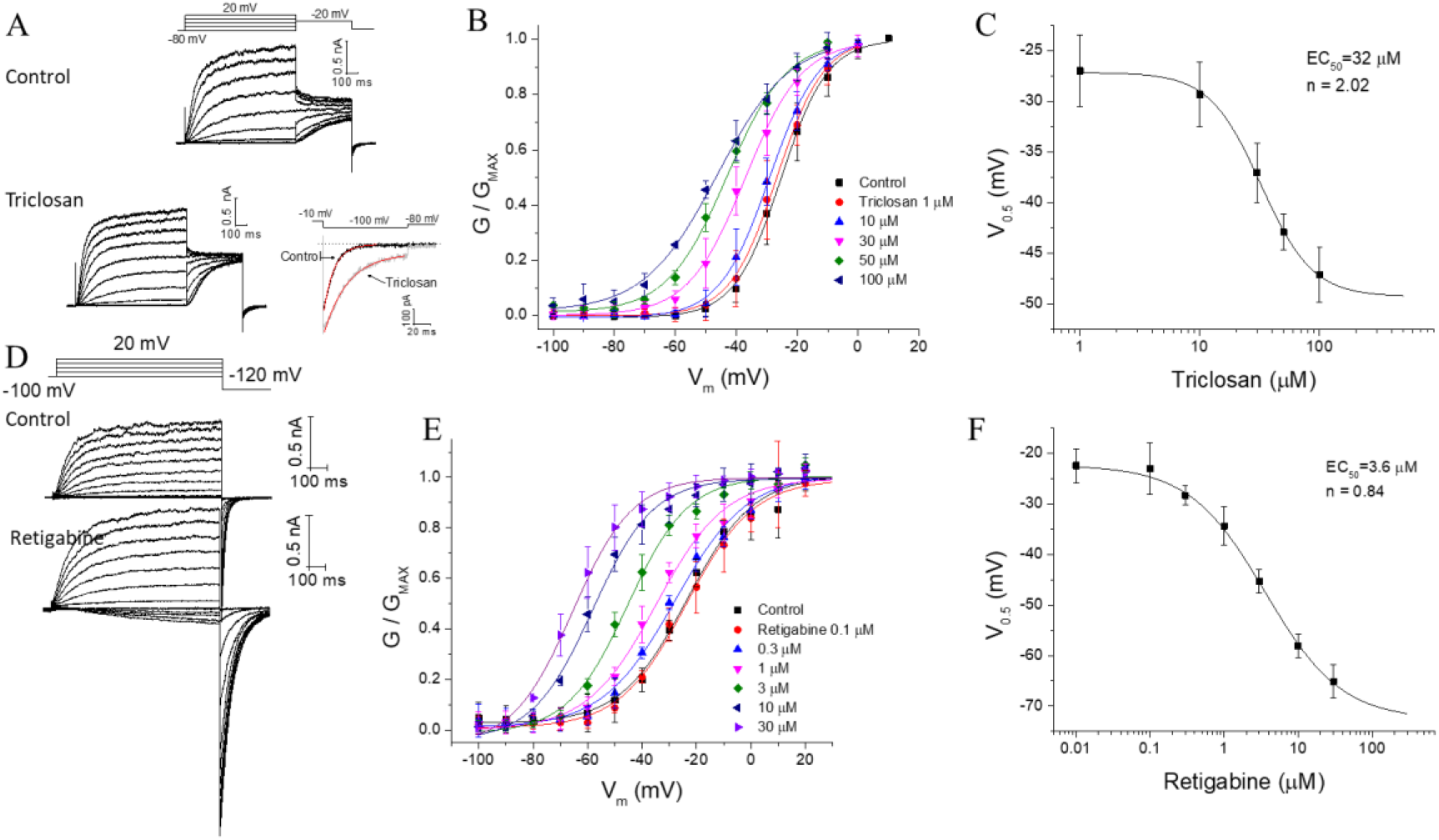
Triclosan activates the KCNQ2/3 heteromers potassium channels. A) representative recordings of KCNQ2/3 channels in control solution and with triclosan 50 μM. B) triclosan induce a shift in the voltage dependence of activation of the channel in a dose dependent manner. The conductance was measured in the tail current at −20 mV. C) dose-response curve with an EC_50_ of 32 μM. D) representative recordings of the KCNQ2/3 with retigabine 10 μM. E) shift in the voltage dependence of the conductance by retigabine. The conductance was measured in the tail current elicited at −120 mV. F) dose-response curve of retigabine. Note that Hill coefficient is different in C and F.

### Triclosan activates KCNQ3 selectively

On homomer KCNQ2 channels (Figure 2A), application of triclosan does not affects the channel voltage dependence. For homomer KCNQ3, we used the KCNQ3-A315T (KCNQ3T) channel, a mutant that increases whole-cell current amplitudes without changing the open probability or PIP_2_ sensibility [29, 30]. Triclosan shifts the open probability of the channel to negative voltages in a dose - dependent manner (Figure 2C and E). The maximum voltage shift is ~80 mV with an EC_50_ = 38 μM and n_H_ = 1.89 (Figure F). To test the possibility that the A315T mutation was responsible for the larger shift compared to KCNQ2/3 channels, the effect of triclosan on KCNQ3 wildtype channel was also evaluated giving the same voltage shift, since the whole cell current on the wildtype channel is small, we continue the experiments on the A315T mutant. To analyze the activation kinetics, a concentration of 35 μM was used and a prepulse to - 120 mV was applied to close the channels (Figure 2G), a slight acceleration of the activation was observed in some cells, however, figure 2H shows that the kinetics is primarily unaffected, and that the increased conductance after triclosan application is mainly due to the shift of the voltage dependence, in fact the voltage shift in figure 2H is the expected shift from the dose-response curve at the concentration used. On the other hand, the inactivation kinetics analyzed at different depolarization steps does not follows solely the voltage shift (see inset on figure 2I), indicating that triclosan affects channel closing. This data indicates that triclosan activates selectively the KCNQ3 channel and that the voltage shift on the activation of KCNQ2/3 channels can be explained solely by this interaction.

**Figure 2.**
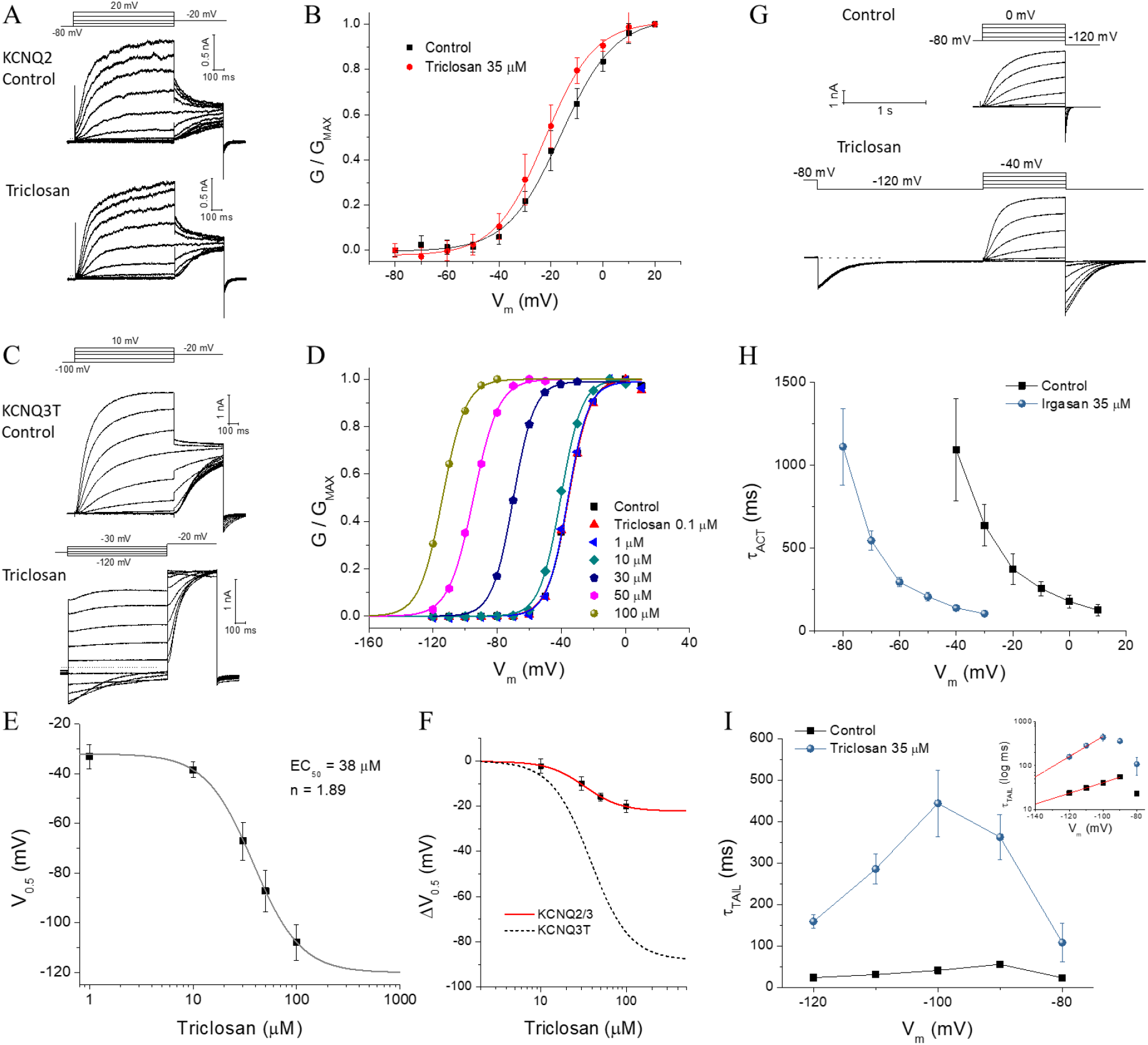
Triclosan activates selectively the KCNQ3 channel. A) representative recordings of KCNQ2 with triclosan 35 μM. Triclosan fails to increase the conductance or shift the voltage dependence of activation (B). C) representative recordings of KCNQ3T in control conditions and with triclosan. D) shift in the voltage dependence of activation the KCNQ3T by increasing concentrations of triclosan. E) dose-response fit gives an EC_50_ and Hill coefficient values similar to KCNQ2/3 channels. F) comparison of the voltage changes in the V0.5 (ΔV_0.5_) caused by triclosan in KCNQ2/3 and KCNQ3T. G) protocol for quantifying the activation time constant in the presence of triclosan, a prepulse to −120 mV was applied to close the channels. The scale applies for both recordings. H and J) comparison of the activation and deactivation time constant in the presence of triclosan. The activation time constant is not affected significantly, instead the deactivation is slower. The inset in (I) is the time constant in logarithmic scale to show the differences more precisely, the red lines are exponential fil to the first part of the data, extrapolated to more negative values. If the time constant is not affected, it would be expected to have similar time constant values ~40 mV apart, as shown for the activation in (H).

### The effect of triclosan is independent of PIP2

It is known that phosphatidylinositol 4,5-bisphosphate (PIP_2_) regulates KCNQ channels activity [31]. We tested if triclosan alters PIP2 binding. We transfected KCNQ3T together with the M1 muscarinic receptor and evaluated triclosan action in the presence of oxotremorine-M. Figure 3A shows that oxotremorine-M (10 μM) inhibits 30% of the current as reported earlier [32], application of triclosan 35 μM still shifts the voltage dependence of activation. The low depression of the KCNQ3 current after M1 receptor stimulation is due to its high PIP2 affinity compared to all other KCNQ channels [33, 34], moreover M1R stimulation decreases only ~80% of PIP2 abundance. To account for these difficulties, we made use of a voltage sensitive phosphatase which can dephosphorylate nearly all PIP2 in the plasma membrane [35]. Activation of *Danio rerio* VSP (Dr-VSP) at +120 mV causes a decay in the KCNQ3T current, upon stepping back to +30 mV, where the Dr-VSP is not activated, the current slowly recovers (Figure 3B and G). This protocol gives an estimate of changes in *k_off_* and *k_on_* of PIP2 from the channel. Triclosan does not significantly alters the decay nor the recovery rates in KCNQ3T, indicating that the *k_off_* and *k_on_* of PIP2 are not influenced by triclosan, however it is worth noting that the decay and recovery measurements are influenced by the rate of dephosphorylation of PIP2 by the VSP at that voltage and the rate of PI(4)P-5 kinase [32, 36], a more sophisticated deconvolution of those rates is beyond the scope of the experiment. To determine the effect of triclosan without PIP2, we measured the tail current kinetics at −90 mV immediately after PIP2 depletion at +120 mV (Figure 3C). We were not able to determine voltage dependent changes with this protocol, instead we evaluated the tail current kinetics as an indicator of triclosan activation. Triclosan slows the tail current kinetics as expected (Figure 3C and D). Furthermore, we tested triclosan on a low PIP2 affinity KCNQ3T mutant, R364A [32]. Surprisingly, triclosan increases five-fold the amplitude of the current, but the voltage shift was the same as in the wildtype channel (Figure 3E and F). In this mutant, the decay of the current due to Dr-VSP activation is faster and the recovery is slower due to the low PIP2 affinity (Figure 3G), triclosan accelerated the recovery rate but the decay was unaffected. We believe that the recovery rate is influenced by the substantial current amplitude enhancement in this mutant. We turned back to the M1 receptor stimulation to deplete PIP2, application of triclosan in these conditions still activates KCNQ3T (R364A). These data suggest that triclosan activation of KCNQ3 is independent of PIP2.

**Figure 3.**
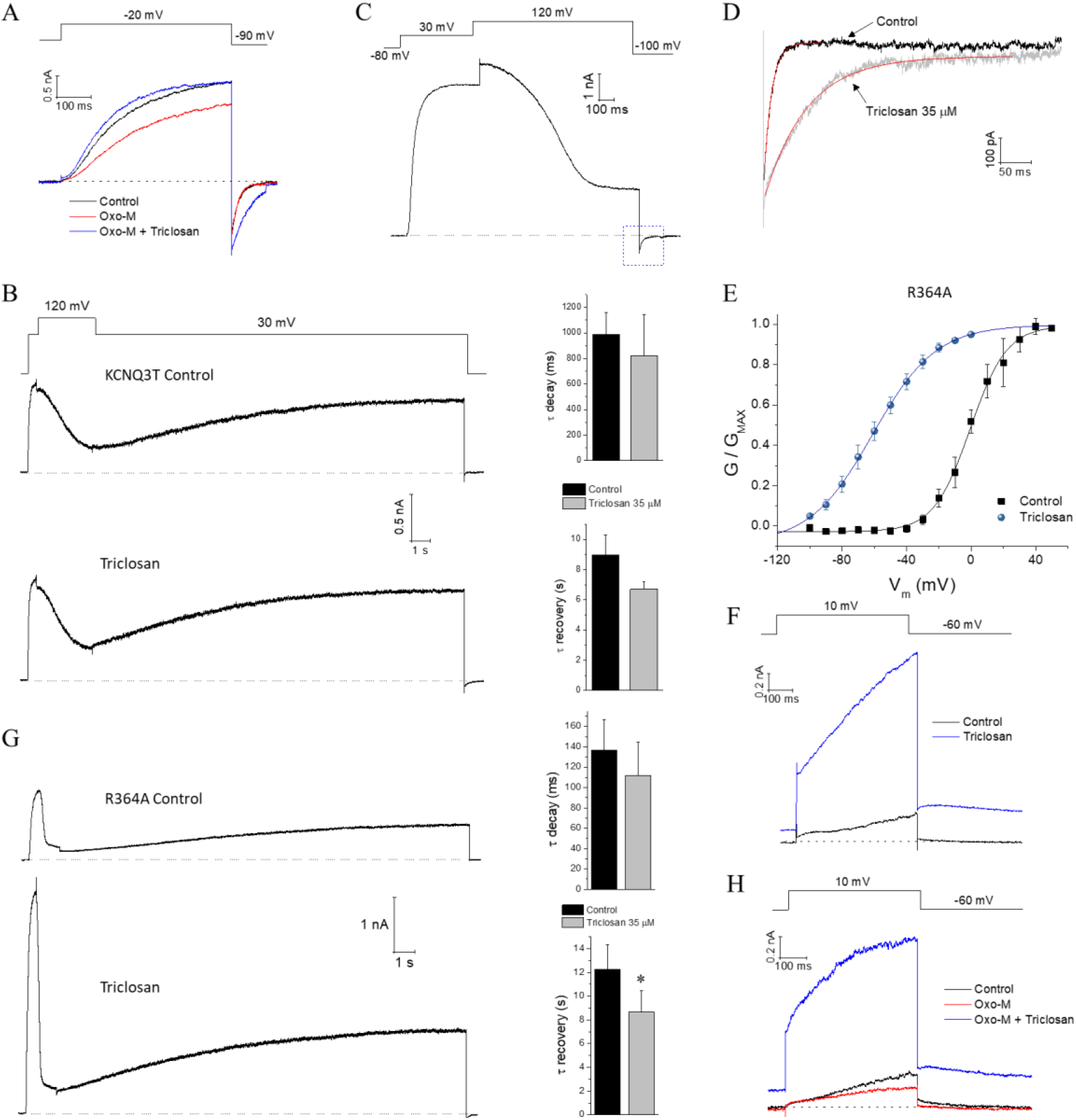
PIP2 does not affect triclosan dependent activation of KCNQ3T. A) cells cotransfected with KCNQ3T and the muscarinic receptor M1. Oxotremorine-M inhibits the current, application of triclosan on this condition still increments the conductance of the channel. B) cells cotransfected with KCNQ3T and Dr-VDP. Activation of Dr-VSP at 120 mV induces a time dependent decay of the current. At 30 mV the Dr-VSP is not activated and the current slowly recovers. The decay and recovery are estimates of the *k_off_* and *k_on_* of PIP2. The parameters are not affected by triclosan. C) tail current elicited at −100 mV after PIP2 depletion by Dr-VSP. D) amplification of the box indicated in (C); the tail current is slower after application of triclosan. E) the low PIP2 affinity mutant KCNQ3T (R364A) is activated by triclosan. The voltage shift in the activation is like the observed in the wildtype channel, however the current increments 5-fold (F). G) Current decay after PIP2 depletion is not affected by triclosan. The recovery is significantly accelerated (* p>0.05). H) M1R activation inhibits the current in the R364A mutant. Triclosan still activates the channel in this condition.

### Putative network of interactions between triclosan and KCNQ3

Triclosan shares some structural similarity to pyridyl benzamide compounds that are characterized as Kv channels activators that bind to the voltage sensing domain [5, 37, 38]. Three residues within the voltage sensor of KCNQ3T where mutated and evaluated for triclosan binding: E160, F167 and S209. Of these, E160A mutant did not produce any visible current, F167A and S209A had the same shift in the voltage dependence as the KCNQ3T channel after application of triclosan (Figure 4A and B). KCNQ2 and KCNQ3 share sequence similarities in the pore region, of particular interest, a tryptophan (W236 in KCNQ2; W265 in KCNQ3) located in S5 has been shown to be critical for retigabine activation. Since triclosan does not affects KCNQ2, the possibility that retigabine and triclosan share the same binding site is minimal. Still, we tested the possibility by making two mutations, W265L and W265A. Both mutants show a leftward shift of the voltage dependence compared to wildtype, −46 ± 3 mV and −73 ± 4 mV, respectively. We switch the bath to a K^+^-free solution to reliably measure the voltage-dependence of activation. Triclosan was still able to activate the channel to the same voltage shift (Figure 4C). A partial dose-response of the W265L is shown in figure 4D, the first derivative of the dose-response data was fitted to a gaussian function (Figure 4E), the peak should give an estimate of the EC_50_ which was close to the EC_50_ determined in the wildtype channel. These results indicate that the binding site for triclosan is different to pyridyl benzamide compounds and retigabine.

**Figure 4.**
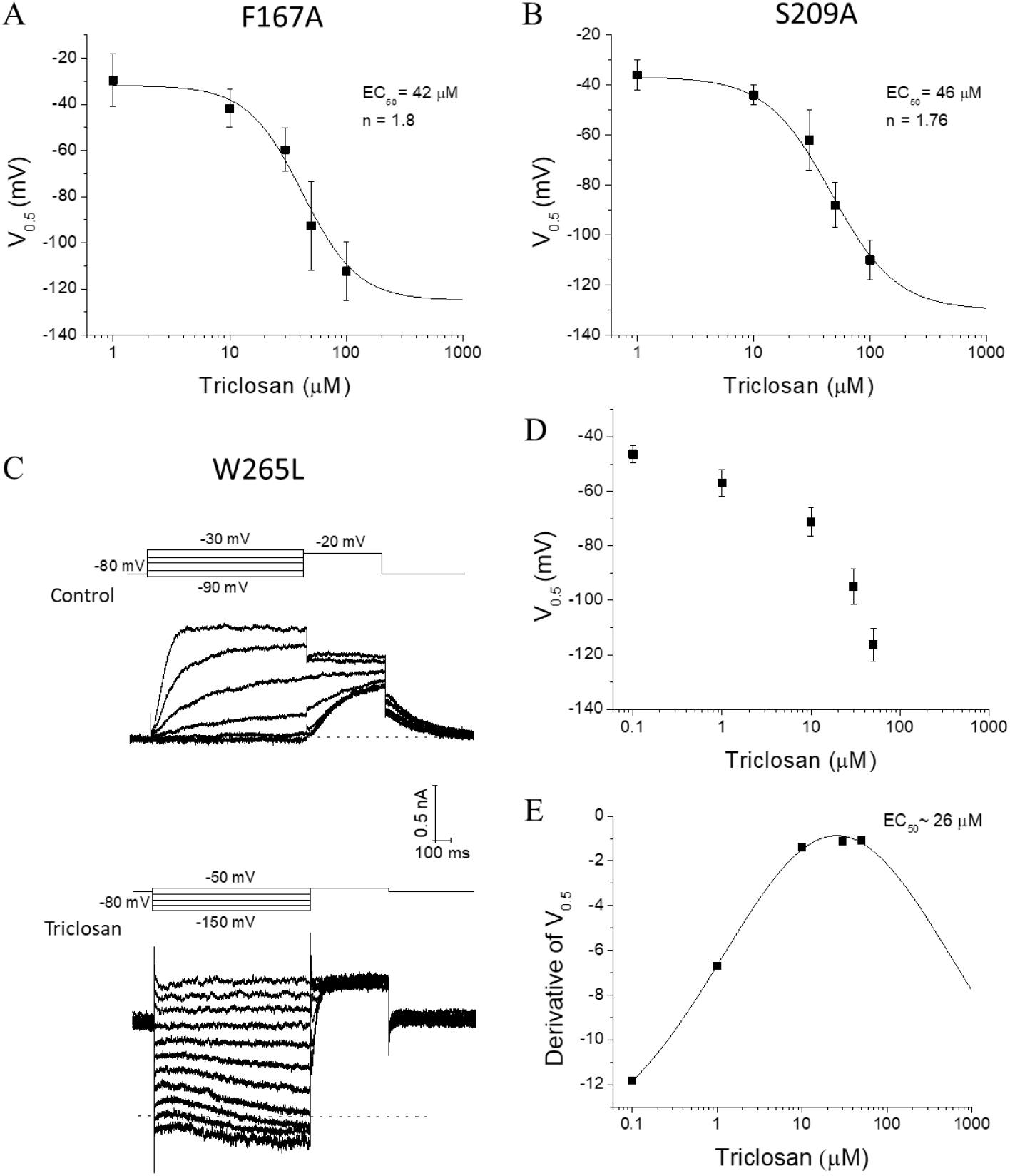
Mutagenesis of possible binding sites for triclosan. A and B) mutations in the voltage sensing domain of KCNQ3T does not affects triclosan activation. F167A and S209A mutant channels have comparable EC_50_ for triclosan. C) mutation of the tryptophan (W265) necessary for retigabine binding does not prevents the activation by triclosan. D) partial dose-response curve for triclosan in the W265L mutant. E) a gaussian fit to the derivative of the data in (D) gives an estimate of the EC_50_. The extracellular solution to record the W265L mutant were K^+^-free.

To model the possible interaction of triclosan with the KCNQ3 channel, we first model the KCNQ3 wildtype channel (Figure 5A and B) based on the KCNQ2 cryo-EM structure as a template (PDBID:7CR0) [22] and perform docking simulations to the most energetically favorable model. The experimental results provided constrictions for the docking simulation. The lowest energy conformation predicts that triclosan binds to the pore turret region (Figure 5C and D) in a network formed by A304, L305, both located in the pore helix; L331 located in the S6 of the adjacent subunit, and L134 located in S1 of the other adjacent subunit. The docking simulation predicts a hydrogen bond between the OH of triclosan and A304.

**Figure 5.**
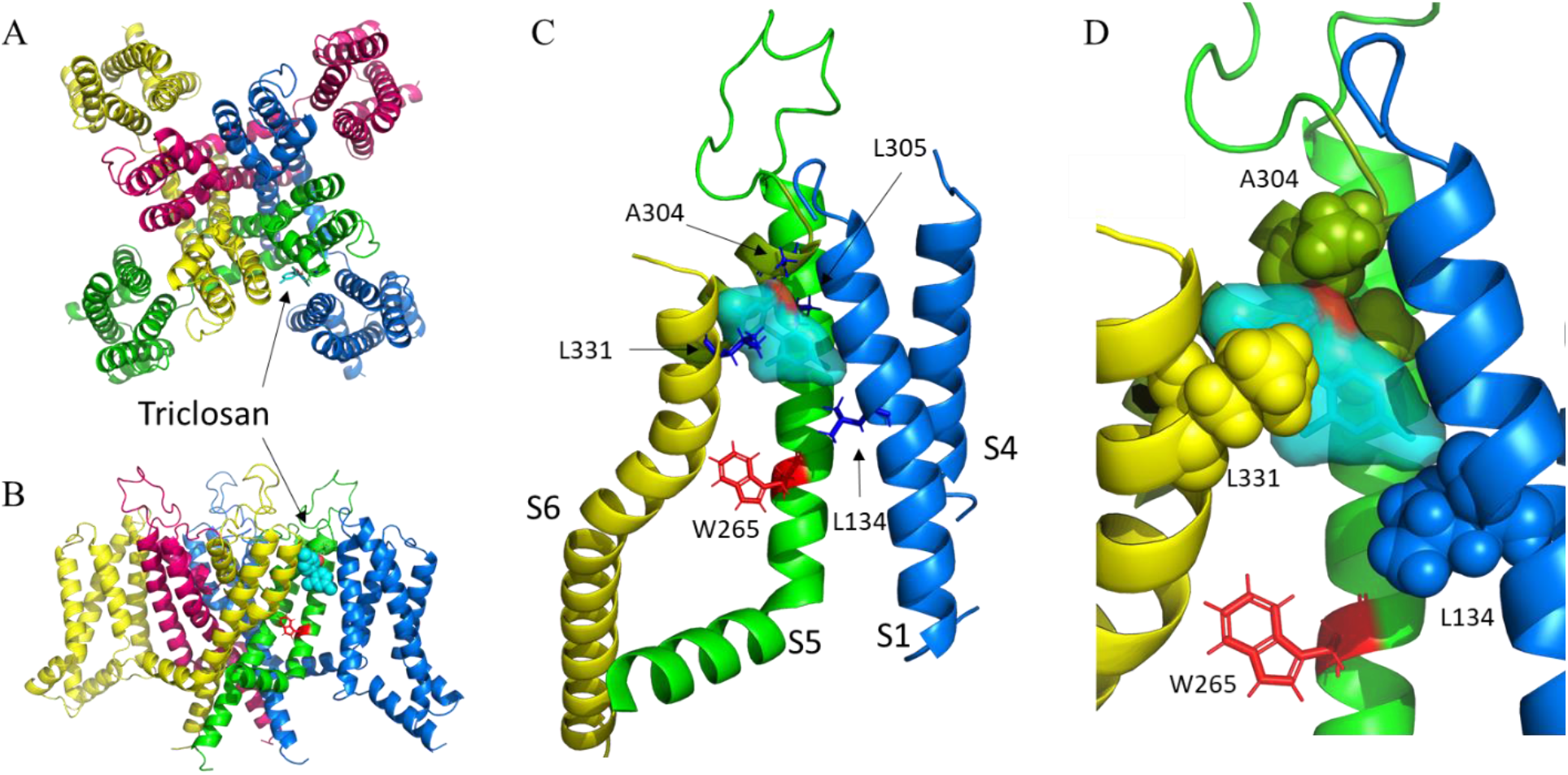
Homology model of the KCNQ3 potassium channel. A) extracellular view. B) side view. Each color represents a different subunit. The voltage sensing domain of the red and green subunits are missing for clarity in (B). Triclosan is represented in sticks in (A) and spheres in (B). C) putative network interactions between triclosan and KCNQ3. The relevant residues are represented in blue sticks. The color of the helix indicates a different subunit. The tryptophan 265, required for retigabine binding is highlighted in red. D) magnification of the image in (C). The relevant residues are now shown in spheres. Triclosan fits almost perfectly into a cavity formed by the indicated residues.

## Discussion

The search for chemical activators of KCNQ channels is still under intensive investigation because of the relevance of these ion channels in treating hyperexcitability-related disorders. The increasing physiological and structural information is helping to unravel new modulation mechanisms and the design of new drugs with increase selectivity and potency. In the present study we identified the widely used bactericide Triclosan as an activator of the KCNQ channels. We show that triclosan activates the heteromeric KCNQ2/3 channels and that the effect is mediated only by the KCNQ3 channel. The kinetics analysis suggests that like retigabine [39], triclosan stabilizes the open state of the channel. Both mutant channels W265L and W265A retain triclosan sensitivity, indicating that the binding site is different from retigabine. Other binding pockets in KCNQ channels have been described for several activators with different subtype selectivity. ICA-27243 shows preferred selectivity for KCNQ2/3 heteromers, but it also modulates KCNQ4 and KCNQ3/5 [40]. The binding site for ICA-27243 and other related compounds like ML213 [41, 42] was found to be in the voltage sensing domain, these compounds increase channel conductance and shift the activation curve to more negative potentials. NSAIDs related compounds have also been described to bind the VSD yielding similar effects on the voltage sensing and channel deactivation [43]. In our experiments we selected three residues that are important for the binding of the above-mentioned compounds, E160, F167 and S209. We were unable to record currents from E160A mutant. F167A and S209A mutants had similar voltage dependent activation as the wildtype channel and triclosan retain the same sensitivity on these mutants, although several other residues need to be tested, these results indicate that the binding site for triclosan differs from these compounds.

The bacteriostatic compound hexachlorophene is a polyphenol structurally related to triclosan that was described as an activator of the KCNQ1/KCNE1 potassium channel, yet it also activates the KCNQ2 and KCNQ4 [44]. This contrast with the selectivity of triclosan that has no effect on KCNQ2 but a profound effect on KCNQ3, we further tested triclosan on KCNQ4 (data not shown) and found that triclosan fail to activate this subtype at 35 μM. It is of interest that hexachlorophene has a slow off rate, evidenced by the washout difficulty after reaching steady state [44]; an observation that contrast with our experiments in which washout of triclosan was completed after ~5 minutes. These results indicate that structural similar compounds have different selectivity of the channel subtypes and that the mechanism of action might also be different.

KCNQ channels are known to be regulated by PIP2, KCNQ3 has an extremely high affinity for PIP2, in fact the open probability is near unity at saturating voltages in contrast to <0.3 for KCNQ2 [33, 34], the lack of sensitivity for triclosan on KCNQ2 opened the possibility for a mechanism dependent on PIP2 for triclosan activation of KCNQ3, however, our results suggest that PIP2 is not necessary for this interaction. Of interest is the five-fold increase on the conductance of the R364A mutant channel, we saw the same result in the H376A mutant channel (only two cells were recorded therefore the data is not shown); both residues are located after the S6 segment, a region that is part of the network of interactions between the channel and PIP2 [32], however, as we show here, triclosan is still able to activate the channel after PIP2 depletion. One possibility is that those mutations are affecting the channel lower gate, or the conformational changes associated with it after triclosan induces changes in the upper half of the pore. The binding of triclosan affects the voltage sensitivity in a similar way as retigabine, although it is not clear how drug binding in the pore region affects the voltage sensing domain, it appears that after retigabine binding, the voltage sensing domain is disturbed, reducing the energy required for the translation between the resting to the activated state [22]. Our observations suggest that a similar mechanism might occur for triclosan given that it can increase the conductance of the R364A and H376A mutants but the voltage shift is unaffected, suggesting that the two effects, voltage shift and conductance increase can be separated. Altogether, our data indicates that triclosan may be a novel class of KCNQ activator. While more drugs with improve features are been discovered, triclosan might be a useful template for the synthesis of new activators of the KCNQ channels that can be used in the therapeutics.

## ACKNOWLEDGEMENTS

The work was supported by grant FORDECYT-PRONACES/1308052/2020 from CONACyT, Mexico.

